# Delirium symptoms are associated with decline in cognitive function between ages 53 to 69: findings from a British birth cohort study

**DOI:** 10.1101/206631

**Authors:** A Tsui, D Kuh, M Richards, D Davis

**Affiliations:** MRC Unit of Lifelong Health and Ageing at UCL, 33 Bedford Place, London WC1B 5JU, UK

**Keywords:** delirium, dementia, cognitive decline, life course

## Abstract

**INTRODUCTION:** Few population studies have investigated whether longitudinal decline after delirium in mid-to-late life might affect specific cognitive domains.

**METHODS:** Participants from a birth cohort completing assessments of search speed, verbal memory and the Addenbrooke’s Cognitive Examination at age 69 were asked about delirium symptoms between ages 60-69. Linear regression models estimated associations between delirium symptoms and cognitive outcomes.

**RESULTS:** Period prevalence of delirium between 60 and 69 was 4% (95% CI 3.2%,4.9%). Self-reported symptoms of delirium over the seventh decade were associated with worse scores in the Addenbrooke’s Cognitive Examination (−1.7 points, 95% CI −3.2, −0.1, p=0.04). In association with delirium symptoms, verbal memory scores were initially lower, with subsequent decline in search speed by age 69. These effects were independent of other Alzheimer’s risk factors.

**DISCUSSION:** Delirium symptoms may be common even at relatively younger ages, and their presence may herald cognitive decline, particularly in search speed, over this time period.

## Introduction

Delirium is a neuropsychiatric syndrome characterized by acute cognitive dysfunction and attentional deficits precipitated by acute illness. Delirium is common, affecting around 25% of older inpatients,[1, 2] and is associated with increased length of hospital admission, risk of institutionalization, and mortality.[3] In a range of settings, delirium has been linked with subsequent cognitive decline and incident dementia.[4–7] The precise mechanisms underlying these relationships are unknown but may be independent of classical Alzheimer’s pathologies such as amyloid beta, hyperphosphorylated tau or APOE-ε4 genotype.[8–10]

While delirium clearly has an impact on long-term cognition, current understanding is incomplete in a number of ways. Firstly, delirium has mainly been investigated in hospital samples,[11–13] and population studies are less common.[14, 15] This is relevant because not all patients with delirium can be assumed to be admitted to hospital; those who are may have more severe or prolonged episodes.[16] Second, there is little knowledge of whether cognitive function after delirium declines in specific domains or whether the observed associations relate to cognition more generally. Thirdly, there are few studies with prospective assessments of cognition, before *and* after delirium, which are necessary in order to quantify prior vulnerability to delirium and its subsequent effects.[6, 14, 17, 18] None of these studies are exclusively in populations younger than age 70. It is likely that delirium and dementia have shared and interacting risk factors, though when these processes start in the life course is unclear.

Our aims were to investigate the association between reported delirium symptoms and change in two key domains of cognition, episodic memory and processing speed between ages 53 and 69 years, using data from a British birth cohort study, accounting for the influences of prior cognitive function as well as early and midlife risk factors for Alzheimer’s disease. We addressed three questions: 1. What is the ten-year period prevalence of reported delirium symptoms in a birth cohort aged 69? 2. Are delirium symptoms subsequently associated with deficits in any particular cognitive domains? 3. Are reported delirium symptoms associated with cognition at age 69, and cognitive decline since midlife independently of other factors known to be associated with Alzheimer’s risk?

## Methods

The MRC National Survey for Health and Development (NSHD) is a British birth cohort study, following a sample of 5362 participants born in March 1946. In 2015, when participants were aged 69 years, 2698 individuals living in England, Scotland and Wales were invited to have a home visit by a research nurse as part of the 24th follow up.[19] The other 2664 participants had either died (n=995), permanently refused participation (n=654), moved abroad (n=583) or were lost and untraced (n=432). Of the 2698 invited, 2148 (80%) completed a home visit and the maximum sample for this analysis is the 2090 who responded to a question about delirium symptoms (97%).

### Cognitive outcomes at age 69

Main outcome variables were verbal memory and visual search speed at age 69. All cognitive assessments were carried out by the research nurse according to the same standardized protocol as at earlier ages.[20] Verbal memory was assessed using a 15-item word learning task, where each word was presented for 2 seconds. The score represents the total number of words correctly recalled over three identical trials (maximum 45). Visual search speed was assessed by crossing out the letters P and W, randomly embedded within a grid of other letters, as quickly and accurately as possible in 1 min. The score represents the total number of letters searched (maximum 600). For the first time, participants were also administered the Addenbrooke’s Cognitive Examination (version III) (ACE-III), which gives scores in five domains: attention & orientation (scored 0-18); verbal fluency (0-14); memory (0-26); language (0-26); and visuospatial function (0-16) (total score 100).

### Delirium symptoms

At age 69, participants reported whether they had experienced an episode of delirium in the previous ten years: *“Please think to a time when you have been unwell, perhaps while in hospital. Sometimes a person’s memory, thinking and concentration can get worse over hours and days due to an illness e.g. infection, operation or due to medications. This is delirium. Since 2006, have you experienced delirium symptoms?”*

### Covariates

We used previous scores of verbal memory and visual search speed at age 53 (in 1999) as a baseline measure predating any delirium from 2006.[20] Other covariates were selected from the data collection at age 60-64,[21] based on factors previously demonstrated to be important in the primary prevention of Alzheimer’s dementia. These were: diabetes,[22] smoking,[23, 24] hypertension,[25] body mass index (BMI),[21] educational attainment[23, 26, 27] and level of physical exercise.[24, 28]

At age 60-64, diabetes was defined as a diagnosis of type 1 or 2 diabetes mellitus by a doctor. Hypertension was defined as a physician-diagnosed hypertension, or regular prescription of an anti-hypertensive, or systolic blood pressure greater than 140mmHg or diastolic blood pressure greater than 90mmHg taken from two measurements. To assess BMI, height and weight were measured by standardized protocols using stadiometers and scales. Smoking status was defined as: current smoker, ex-smoker, or lifelong non-smoker, corroborated with reports at earlier ages. Participants were asked how many times in the last four weeks they had taken part in sports or vigorous activities, classified as inactive (no episodes), less active (1-4 exercise episodes per month) and more active (five or more exercise episodes per month). Educational qualifications by age 26 were categorized into no educational qualifications or less than ordinary ‘O’ level; ‘O’ levels; advanced ‘A’ level and higher.

### Statistical Analysis

First, using the maximum samples, the cognitive test scores (verbal memory, search speed, ACE-III and its subdomains) at ages 69 and all covariates were assessed in respect to reported delirium symptoms using chi squared tests and t-tests as appropriate. Second, for participants with complete data for visual search speed and verbal memory at 53 and 69, we used linear regression to estimate the associations between delirium symptoms and individual cognitive domain at age 69, adjusting only for sex and the corresponding cognitive measure at age 53. We then used all other covariates in a multivariable model. We tested for interactions between delirium and each covariate in the fully adjusted models. Finally, we tested the associations between delirium symptoms and ACE-III measures (total scores), in order to derive a comparable estimate of effect size on a clinically relevant scale. Residuals for all regression models were checked for heteroscedasticity. Stata 14.1 (StataCorp, Texas) was used for all analyses.

## Results

Of 2090 participants responding to the delirium question, 83 reported symptoms between ages of 60 to 69, thus a ten-period period prevalence of 4.0% (95% CI 3.2% to 4.9%). No sex differences were observed (p=0.59). Complete ACE-III data were available for 1729 participants (81% of home visits). Participant characteristics for maximum samples by reported delirium symptoms are shown in Table 1.

**Table 1.** Patient characteristics of study participants with and without symptoms of delirium between ages 60-69

|  | **n** | **Delirium symptoms** | **No delirium symptoms** | **P** |
| --- | --- | --- | --- | --- |
| Visual search speed (letters, SD) |  |  |  |  |
| Age 69 | 2053 | 237 (76) | 263 (74) | <0.01 |
| Age 53 | 1968 | 280 (76) | 284 (73) | 0.67 |
| Verbal memory (words, SD) |  |  |  |  |
| Age 69 | 2018 | 21 (6) | 22 (6) | 0.14 |
| Age 53 | 1955 | 23 (7) | 25 (6) | 0.04 |
| Hypertension at age 60-64 | 1829 | 22% | 17% | 0.25 |
| Diabetes by 60-64 | 1956 | 15% | 11% | 0.27 |
| Smoking status at 60-64 | 1808 |  |  | 0.63 |
| *Current* |  | 13% | 10% |  |
| *Ex* |  | 59% | 57% |  |
| *Never* |  | 28% | 33% |  |
| Level of activity age at 60-64 | 1923 |  |  | 0.39 |
| *Inactive* |  | 66% | 58% |  |
| *Less active (1-4/wk)* |  | 11% | 13% |  |
| *More active (>5/wk)* |  | 23% | 29% |  |
| Educational Attainment by 26 | 2090 |  |  | 0.63 |
| <O Levels |  | 37% | 40% |  |
| O Levels |  | 20% | 16% |  |
| >=A Levels |  | 44% | 45% |  |
| BMI aged 60-64 | 1826 | 28.7 (4.5) | 27.6 (4.8) | 0.19 |

*Chi square p value for categorical variables, t-test for continuous variables*

At age 69, the mean search speed was lower for those who reported a history of delirium symptoms than those reporting none (mean difference -26 letters; p<0.01) (Table 1). There was little difference in verbal memory by reported delirium symptoms (mean difference -1 words; p=0.14). At age 53, no difference in search speed was found between those with and without subsequent delirium symptoms (mean difference −4 letters; p=0.67). In contrast, mean verbal memory was lower for those with subsequent delirium symptoms (mean difference −1.6 words; p=0.04). No associations were found between delirium symptoms aged 60-69 and hypertension, diabetes, smoking status, level of activity, BMI and educational attainment (Table 1). Higher levels of exercise at age 60-64 were associated with faster search speed and higher memory at 69, while greater educational attainment and female sex were associated with higher verbal memory at 69 (Supplementary Table 1).

Delirium symptoms were associated with lower total ACE-III scores (89.5 points versus 91.6 points, p<0.01) (Table 2). When examining subscores in the cognitive domains, delirium symptoms were specifically associated with worse performance in memory and visuospatial function (Table 2).

**Table 2.** The relationship between history of delirium symptoms and the Addenbrooke’s Cognitive Assessment (Version 3) at age 69.

| **Mean ACE-II Score** | **Delirium symptoms** | **No delirium symptoms** | **P** |
| --- | --- | --- | --- |
| Total /100 | 89.5 | 91.6 | <0.01 |
| Attention and orientation / 18 | 16.7 | 16.7 | 1.00 |
| Verbal fluency / 14 | 10.5 | 11 | 0.55 |
| Memory /26 | 22.5 | 23.4 | 0.01 |
| Language / 26 | 24.9 | 25.3 | 0.05 |
| Visuospatial /16 | 14.9 | 15.1 | <0.01 |

The estimated association between delirium symptoms and lower search speed did not attenuate after sequentially adjusting for sex, prior visual search speed at age 53, and the full set of covariates (Table 3). In this fully adjusted model, those reporting delirium symptoms searched 31 fewer letters (95% CI −41 to −12, p<0.01) (Table 3). There was no association between reported symptoms of delirium and verbal memory performance at age 69, adjusting for covariates using the same multivariable model (p=0.56). For the ACE-III, delirium symptoms were associated with −2.4 points (95% CI −4.1 to −0.7, p=0.01) after adjusting for search speed at age 53. This association remained, even after accounting for other covariates (−1.7 points, 95% CI −3.2 to −0.1, p=0.04) (Table 3).

**Table 3:** Differences in a) visual search speed and b) verbal memory c) ACE-III at 69, by history of delirium symptoms at age 60-69.

|  | n | Adjusted for sex |  |  | + Adjusted for prior cognition* |  |  | + Adjusted for all covariates† |  |  |
| --- | --- | --- | --- | --- | --- | --- | --- | --- | --- | --- |
| | | $\beta$ | 95% CI | P | $\beta$ | 95% CI | P | $\beta$ | 95% CI | P |
| Visual search | 1529 | -32 | (-53, -10) | <0.01 | -33 | (-51, -14) | <0.01 | -31 | (-49, -12) | <0.01 |
| Verbal Memory | 1511 | -0.9 | (-2.6, 0.8) | 0.30 | -0.6 | (-1.9, 0.7) | 0.38 | -0.4 | (-1.7, 0.9) | 0.56 |
| ACE-III | 1323 | -2.4 | (-4.2, -0.7) | 0.01 | -2.4 | (-4.1, -0.7) | 0.01 | -1.7 | (-3.2, -0.1) | 0.04 |

\*ACE-III adjusted for search speed at 53.
† hypertension, diabetes, BMI, smoking status, physical activity at age 60-64, education by age 26
Visual search = letters searched; verbal memory = words recalled; ACE-III Addenbrooke's Cognitive Examination (3<sup>rd</sup> version), total score out of 100.

## Discussion

In a large representative British population cohort at age 69, the ten-year period prevalence of self-reported delirium symptoms was 4%. Self-reported symptoms of delirium were associated with lower search speed at age 69 and faster decline in this domain since age 53. Similar longitudinal associations were not seen for verbal memory as measured by a word list task, though cross-sectional differences were apparent for the ACE-III memory and visuospatial subscores. Taken together, our findings suggest that these self-reported symptoms of delirium appear to contribute to decline in different cognitive domains, and across ages younger than previously described.

The major strength of this study derives from its prospective nature, allowing for the assessment of the independent associations between prior cognitive test scores and selfreported symptoms of delirium on cognitive outcomes. The main weakness of this study was the reliance on self-report when defining delirium in the community. Self-reported delirium may have been limited by recall bias or inability to recognize delirium from the nurse description. Delirium recall, at least in hospital samples, will be less reliable in those with amnestic symptoms as part of their delirium, but also in those with more perceptual disturbance and indeed those with more severe delirium.[29] Nonetheless, in the absence of an established instrument to identify delirium in community samples, it would appear that positive responses to the questionnaire as answered in our study remains informative in respect of the association between delirium symptoms and worsening cognition. Another notable limitation is that, delirium symptoms were recalled over between 60-69 years though cognition was assessed between ages 53 and 69; this discrepancy means that cognitive decline may have taken place in the 7 years after the first cognitive assessment, before any self-reported symptoms of delirium. This is less likely because we have shown that the rate of cognitive decline was faster in the latter period.[30] In common with other cohort studies, our findings may be subject to residual confounding and may be specific to the population and era under study.

The influence of delirium on later-life cognitive impairment had been reported in older adults.[4, 6, 14, 17] Our study is the first to describe and characterize the associations between self-reported symptoms of delirium in the community and later-life cognition at relatively younger ages of 60 to 69, though there have been major studies in younger perioperative[7] and critical care[31] cohorts. Also, while studies had reported associations between delirium and global cognition scores such as the information-memory-concentration subtest of the Blessed Dementia Rating Scale[4] and the Mini-Mental State Examination,[5] our study has some domain-specific findings.

Domain-specific relationships between self-reported symptoms of delirium and cognition appear to vary over the life course. Verbal memory scores were already lower at baseline in those reporting delirium symptoms later in life; while decline in visual search speed only appeared after the self-reported symptoms of delirium. At age 69, the ACE-III scores indicated particular deficits in episodic memory (slightly broader in scope than our longitudinal word-learning verbal memory task) and visuospatial functions. Insofar as the relationship between delirium and dementia is bidirectional, lower verbal memory at baseline may suggest an increased susceptibility to delirium episodes during acute illness, while declining search speed and visuospatial functions are perhaps adversely affected by processes related to delirium. Whether particular structural or functional neural networks are particularly vulnerable to the development and recovery of delirium is unclear. However, demonstrating cognitive decline after delirium symptoms, after accounting for Alzheimer’s risk factors, strengthens the emerging idea that delirium affects cognition via processes independent of classical dementia mechanisms.[10]

Overall, this analysis suggests that episodes of self-reported symptoms of delirium, as early as in the seventh decade, are associated with greater decline in processing speed, and may differentially affect other cognitive domains. These relationships require further replication in other samples, both observational and experimental. Our findings highlight the importance for clinicians to assess cognitive function in patients who at highest risk of delirium, and subsequently preventing and managing delirium to minimize adverse cognitive sequelae in later life.

## Declarations

### Ethics approval and consent to participate

Ethical approval was obtained from the Multicentre Research Ethics Committee (for data collections up to 2010) and Queen Square Research Ethics Committee (14/LO/1073) and the Scotland A Research Ethics Committee (14/SS/1009) for data collections between 2014 and 2015.

### Availability of data and material

Bona fide researchers can apply to access the NSHD data via a standard application procedure (further details available at: http://www.nshd.mrc.ac.uk/data.aspx).

### Competing interests

The authors declare that they have no competing interests.

### Funding

The NSHD, DK, MR and AT are supported by core funding and grant funding (programme codes: MC_UU_12019/1, MC_UU_12019/3) from the UK Medical Research Council. DD is funded through a Wellcome Trust Intermediate Clinical Fellowship (WT107467). The authors declare no conflicts of interest. The study sponsors were not involved in the study design; in the collection, analysis, interpretation of data; in the writing of the report; or in the decision to submit the article for publication. The researchers worked independently from funders. All authors, had full access to all of the data (including statistical reports and tables) in the study and can take responsibility for the integrity of the data and the accuracy of the data analysis.

### Authors’ contributions

AT designed the study, analysed the data, drafted and revised the paper. DD designed the study and revised the draft paper. He is the guarantor. DK and MR advised on the statistical analysis plan and revised the paper.

## Acknowledgements

The authors thank all study members of NSHD and NSHD scientific and data collection teams.

